# Genome-wide interologous interactome map (TeaGPIN) of *Camellia sinensis*

**DOI:** 10.1101/696062

**Authors:** Gagandeep Singh, Vikram Singh, Vikram Singh

**Affiliations:** Centre for Computational Biology and Bioinformatics, School of Life Sciences, Central University of Himachal Pradesh, Dharamshala, India-176206

**Keywords:** *Camellia sinensis*, PPI network, Interologous, Tea genome, Pathways, Transcription factors (TFs)

## Abstract

Tea, prepared from the young leaves of *Camellia sinensis*, is a non-alcoholic beverage globally consumed due to its antioxidant properties, strong taste and aroma. Although, the genomic data of this medicinally and commercially important plant is available, studies related to its sub-cellular interactomic maps are less explored. In this work, we propose a genome-wide interologous protein-protein interaction (PPI) network of tea, termed as TeaGPIN, consisting of 12,033 nodes and 216,107 interactions, developed using draft genome of tea and known PPIs exhaustively collected from 49 template plants. TeaGPIN interactions are prioritized using domain-domain interactions along with the interolog information. A high-confidence TeaGPIN consisting of 5,983 nodes and 58,867 edges is reported and its interactions are further evaluated using protein co-localization similarities. Based on three network centralities (degree, betweenness and eigenvector), 1,302 key proteins are reported in tea to have *p*-value < 0.01 by comparing the TeaGPIN with 10,000 realizations of Erdős-Rényi and Barabási-Albert based corresponding random network models. Functional content of TeaGPIN is assessed using KEGG and GO annotations and its modular architecture is explored. Network based characterization is carried-out on the transcription factors, and proteins involved flavonoid biosynthesis and photosynthesis pathways to find novel candidates involved in various regulatory processes. We believe the proposed TeaGPIN will impart useful insights in understanding various mechanisms related to growth and development as well as defence against biotic and abiotic perturbations.

## Introduction

Topological structure of biological systems is highly complex at every scale of organisation. Several signalling and transcriptional regulatory mechanisms that are supposed to have been evolved to counter various forms of environmental or pathogenic stresses are key contributors of such organizational complexity [1,2]. Network biology finds its crucial role in analysing the functional aspects of a system by explicating several modulatory maps to understand how various subsystems are interwoven together [3]. One of the main reasons of complex behaviour in an organism is the continuous regulation of cellular dynamics that results due to interactions among various biomolecules. Proteins interact with each other through various processes like transduction, translation, cell signalling and metabolism, etc.; the detailed map of exhaustive interactions among all the constituent proteins of an organism is termed as the protein-protein interaction network (PPI) [4,5]. Due to the advancement in high throughput and computational methods, it is now possible for researchers to study and analyse all possible aspects of protein-protein interactions in a cell which in turn enhances the knowledgebase related to dynamics of cellular behaviour [6]. PPIs have major roles in understanding the crucial insights of almost all biological processes and are also helpful in providing new functional sites for unknown pathways and functions [7]. Experimentally determined PPIs have been reported in literature for several organisms including *Homo sapiens* [8]., *Saccharomyces cerevisiae* [9,10], *Drosophila melanogaster* [11] *etc.*, however, these are far from being complete as determining a complete set of PPIs is highly time and cost intensive task. Although, experimental identification of PPIs is always preferred, however, due to its cost and time intensive requirements *in-silico* alternatives are being sought. One such alternative is interolog method where interactions are transferred from other plant species with known PPI information to plants for which PPI data is lacking by exploiting the principle of homology. So, a link is placed between two proteins in query network only if there exists a link between their orthologous proteins in any of the template network. Interologous based PPI networks have been widely studied in various crops including *Arabidopsis thaliana* [12], *Oryza sativa* [13], *Zea mayz* [14], etc. to explain the complex biological problems.

In the class of non-alcoholic beverages, after water, tea (*Camellia sinensis*) is globally the most consumed one mainly due to presence of various antioxidants, pleasant flavours and aroma. It is a valuable source of secondary metabolites such as alkaloids, flavonoids, theanine and other polysaccharides, thus contributing to various health benefits [15]. Tea is a widely researched plant species and is being studied in detail due to its medicinal properties. This crop suffers from various kinds of stresses (Biotic and abiotic), and therefore, to understand the various mechanisms at molecular level to encounter the stress conditions, an interactomic map of whole proteome can play a significant role. Recently, we have developed an interologous method based PPI network of the *Camellia sinensis* leaf using its transcriptomic data from six stress conditions [16]. In order to explore the global understanding of protein interactions in this species, in this work, we are proposing its genome wide PPI network (TeaGPIN). In this study, whole proteomic data of *Camellia sinensis* was incorporated in order to construct an interolog based PPI network of tea, and key regulatory candidates are reported by developing a large number of corresponding random networks and comparing three network properties namely betweenness, degree and eigenvector centralities. Among the key proteins, transcription factors belonging to all the 58 categories are predicted that may be playing crucial roles in various metabolic, signaling and system level pathway regulation. Additionally, two categories of transcription factors (SNF2, responsible for chromosome stability, and C3H, involved in RNA pre-processing) are also explored.

We have also identified proteins potentially involved in flavonoid biosynthesis pathways and studied their sub-network, as flavonoids are crucial secondary metabolites having roles in growth and development of tea plant as well as having health promoting antioxidant properties [17]. Furthermore, sub-network of proteins involved in photosynthetic pathway is also developed and as a specific case, interactions of two important proteins related to photosynthesis (plastocyanin and ferredoxin NADP+ reductase) are investigated in detail as these proteins have important role in electron transport and reductase activity. We propose that these key candidates may be selected for the improvement of crop quality and production through genetic engineering or molecular breeding methods.

## 2. Materials and Methods

### 2.1. Data collection and identification of orthologs

Due to immense medicinal importance, tea has gained unprecedented attention from researchers across various disciplines in current times. Recently two genomes of *camellia sinensis* have been sequenced and are available in public domain [18,19]. As the genome sequence reported by Wei et al., 2018 is available with draft annotation with coding part of genome in the form of translated proteins; this tea proteome was selected for further investigations. A total of 49 plant species having protein interactions in STRING, as available on January 2019, were selected for interolog identification [20]. The proteome sequences of selected plants were obtained from UniProt database (http://www.uniprot.org/) and subjected to BLASTp against tea proteome with e-value ≤ 10^−10^ in order to first identify orthologous proteins [16].

### 2.2 Construction of genome wide protein-protein interaction (PPI) network of tea (TeaGPIN)

All the protein-protein interactions with confidence score ≥ 0.7 in the selected 49 plants were extracted from STRING database [20]. Further, based on interologous network construction approach, the orthologous pairs of tea proteins found in each of the selected plant species were used to construct an exhaustive PPI network of *Camellia sinensis* and this network is termed as TeaGPIN. An interaction between any pair of tea proteins is included in TeaGPIN, if any of the 49 template plants contains that particular interaction [5]. The final constructed TeaGPIN was visualized and analyzed using the Cytoscape v3.3.0 [21]. Once the network was constructed, its giant component was considered for further analysis and interpretation of the network.

### 2.3. Ranking of predicted protein-protein interactions to construct a high confidence TeaGPIN (hc-TeaGPIN)

Proteins interact through the specific domains, thus identifying interacting domain pairs in interologs of the developed network should enhance the confidence of predictions. Motivated by this hypothesis, domains of network proteins were predicted from Pfam database [22] and information about the interactions were extracted from DOMINE database [23]. DOMINE database is a collection of 6,634 experimentally known domain interactions and 21,620 computationally predicted domain pairs exhaustively compiled from 13 different resources. Further, each PPI was ranked by associating a confidence score (*S*_*total*_) consisting of two components, *namely*, interolog score (*S*_*ilog*_) and domain-domain interaction (DDI) score [24]. DDI score is further comprised of two terms; domain interaction propensity (*S*_*dip*_) and correction for interacting-domains enrichment (*S*_*cide*_). *S*_*ilog*_ is the number of template plants from which the interaction under consideration is inferred. *S*_*dip*_ is the ratio of interacting domain pairs reported between two proteins involved in a PPI to the total possible interaction pairs between domains present in these two proteins and is called the propensity of domain interaction. Additionally, to account for domain interaction enrichment bias resulted due to the increase in number of domains present in proteins involved in an interaction; a correction factor called (*S*_*cide*_) was introduced [24].

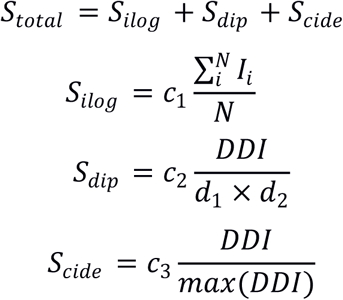

where, *N* represents the total number of template plants used for inferring the interologs and *I*_*i*_ is equal to 1 if an orthologous interaction is transferred from a template plant, and 0 otherwise. *d*_1_ and *d*_2_ represent the number of domains present in two proteins involved in an interaction, while *DDI* is the number of reported domain-domain interactions in DOMINE database among all the possible DDIs (*d*_1_ × *d*_2_) for the respective PPI pair. Constants *c*_1_, *c*_2_ and *c*_3_ are the scaling factors having values 0.5, 0.4 and 0.1 respectively. The confidence scores, thus obtained for each PPI, were then normalized using min-max scaling. All the interologs with normalized *S*_*total*_ ≥ 0.1 were considered as high confidence interologs (HCI).

### 2.4. Protein colocalization similarity (PCS) based reliability assessment of hc-TeaGPIN

PCS method is based on the idea that proteins performing similar functions or involved in common pathways should reside in the same sub-cellular compartments [24]. Sub-cellular locations for each interolog of TeaGPIN was predicted using CELLO tool [25] and the number of interologs having same sub-cellular locations were recorded for the respective locations to develop a distribution. This distribution was compared with the distribution of average of 10,000 ER type random models.

### 2.5. Topological analysis of TeaGPIN

#### 2.5.1. Centrality measures

Giant component of the TeaGPIN was selected for the topological analysis. To identify the key nodes, we have used three network centralities in this study, *namely*, degree, betweenness and eigenvector. For a graph *G* (‖, *E*) with *V* nodes and *E* edges, degree centrality (*C*_*d*_) of a node *v* gives the number of direct links it has with its neighbours. Higher the degree of a node, greater is its capability to control the network [26]. Betweenness centrality (*C*_*b*_) of a node is a measure that computes the ability of a node to act as a bridge between any two nodes [27]. For any two nodes *i, j* | *v* ∈ *V* of *G*, if *σ*_*i,j*_ are the total number paths existing between *i, j* and *σ*_*i,j*_ (*v*) are the paths which are going through node *v*, then the *C*_*b*_ for node *v* is computed as 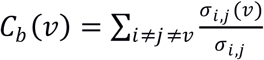. Eigenvector centrality (*C*_*ev*_) of any node is an improved version of degree centrality where alongside considering the degree of an individual node, degree of its first neighbours is also taken into account while ranking that node [28].

#### 2.5.2. Identification of statistically significant proteins in TeaGPIN

*G*_*n,m*_ type Erdős–Rényi (ER) random models preserving order and size of TeaGPIN [29] along with scale free (SF) models based on an extended Barabási–Albert algorithm in which order is preserved and edges differ only within a range of 1% from the original TeaGPIN were used to identify key proteins. For SF models an initial set of *i* isolated nodes was selected in a manner so that *i* ≤ *k* ≤ (*i* + 1), where 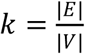. Then, each subsequent vertex was introduced to the initial set with *i* or *i* + 1 number of new edges each having probabilities (*i* + 1 − *k*) and (*k* − 1) respectively [30,31]. Then, 10,000 random realizations of both ER and SF type models were generated. Above mentioned three network centrality metrics were computed for TeaGPIN and for each of the realization in the both sets of random networks. Further, statistically significant key nodes were identified by applying *Z*-statistics that is defined as 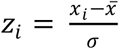.

### 2.6. Functional analysis of TeaGPIN

Sequences corresponding to network nodes were extracted and Gene Ontology (GO) annotation of the proteins was performed by mapping the proteins to TAIR database [32] and AgriGO database [33] Cellular components (CC), biological processes (BP) and molecular functions (MF) were computed using WEGO tool [34]. To find the pathways regulated by the proteins participating in TeaGPIN, KEGG database [35] was used. Furthermore, to identify the transcription factors (TFs), data of all 58 categories of TFs was downloaded from plant transcription factor database [36] and the sequences of each category were aligned using standalone Clustal Omega tool [37]. HMMER was used to build the hidden Markov models (HMM) of each transcription factor category [38]. Network proteins were searched against HMMs of each category of plant transcription factors to find transcription factor activity. Moreover, iTAK [39] tool was also used to identify the transcription factors coding proteins. Additionally, modular architecture of TeaGPIN was explored, as several biological processes are carried out in the cell through interactions among some specified group of proteins. For that, MCODE assisted clustering was done to identify the functional modules in the TeaGPIN [40]. Work-flow chart of the full methodology employed in this work is given in the Figure 1.

**Figure 1.**
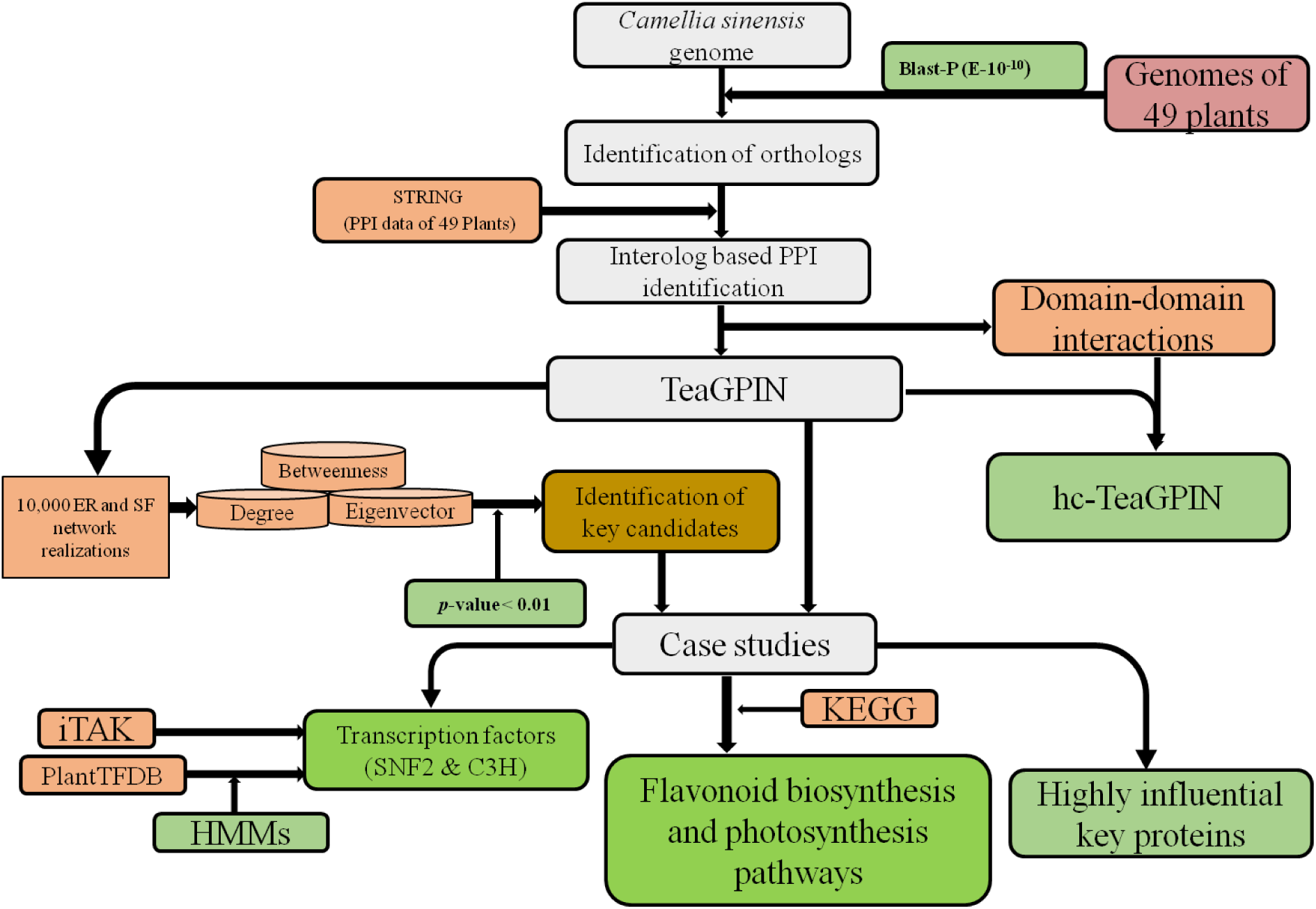
Overall work flow chart involving all the steps in the construction and analysis of TeaGPIN.

## 3. Results and discussions

### 3.1. Construction and analysis of TeaGPIN

The coding part of the publicly available tea genome in the form of 33,932 translated proteins is selected for further analysis [18]. Protein sequences of 49 plant genomes with protein-protein interaction data available in STRING database were selected as templates. The numbers of interacting orthologous pairs (interologs) of tea proteins obtained from 36 of the template proteomes are enlisted in Table 1, from other proteomes no interolog was obtained. All interologs were further selected to generate interolog based PPI network by adding unique interactions from each template. A total of 12,033 nodes having 216,107 interactions could be successfully predicted using the described interolog based method and the resulting network is termed as TeaGPIN as shown in Figure 2A. Degree distribution of the obtained TeaGPIN follows power law with an exponent −1.47, as shown in Figure 2C. (All interactions are given in Supplementary table S1).

**Table 1.**
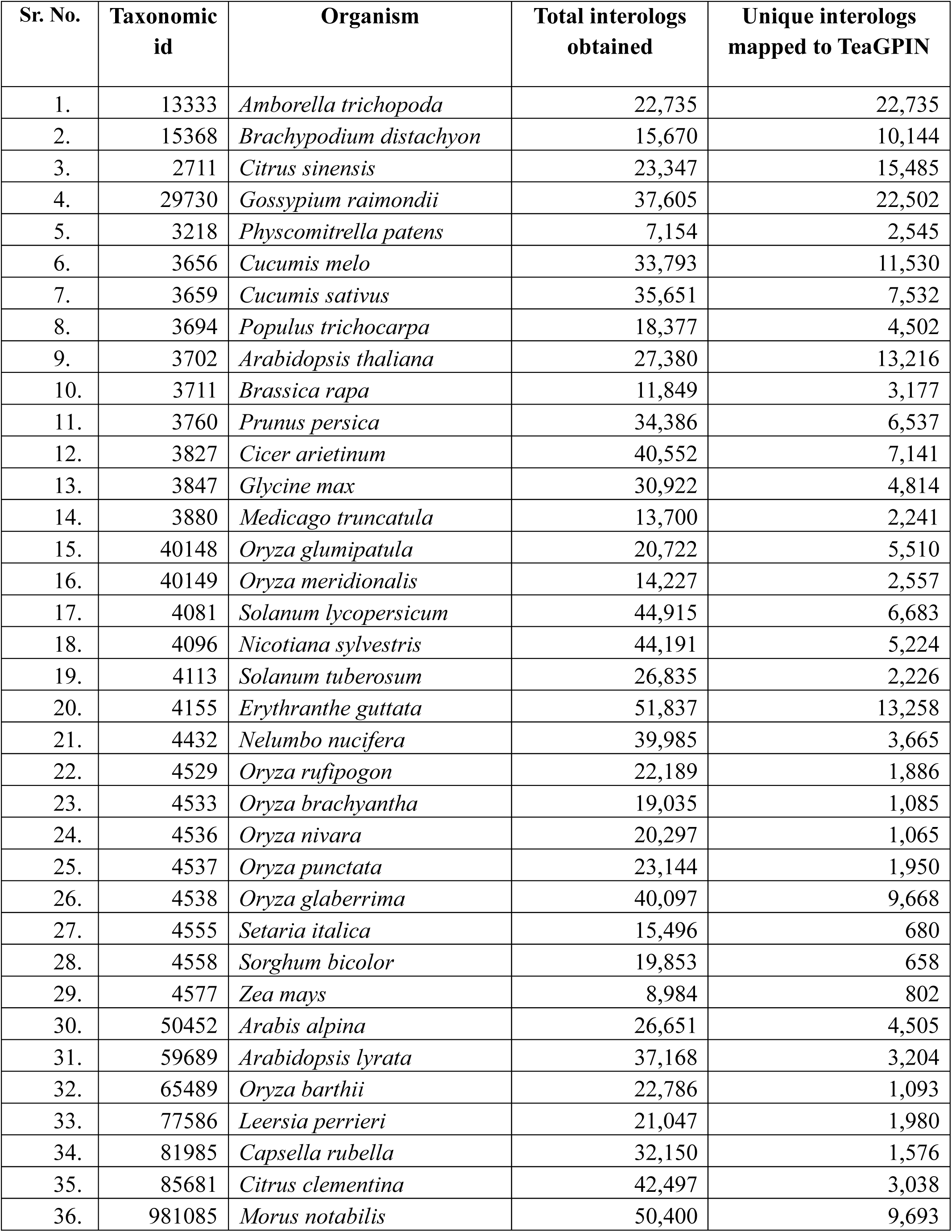
Details of TeaGPIN interologs obtained from 36 template plants.

**Figure 2.**
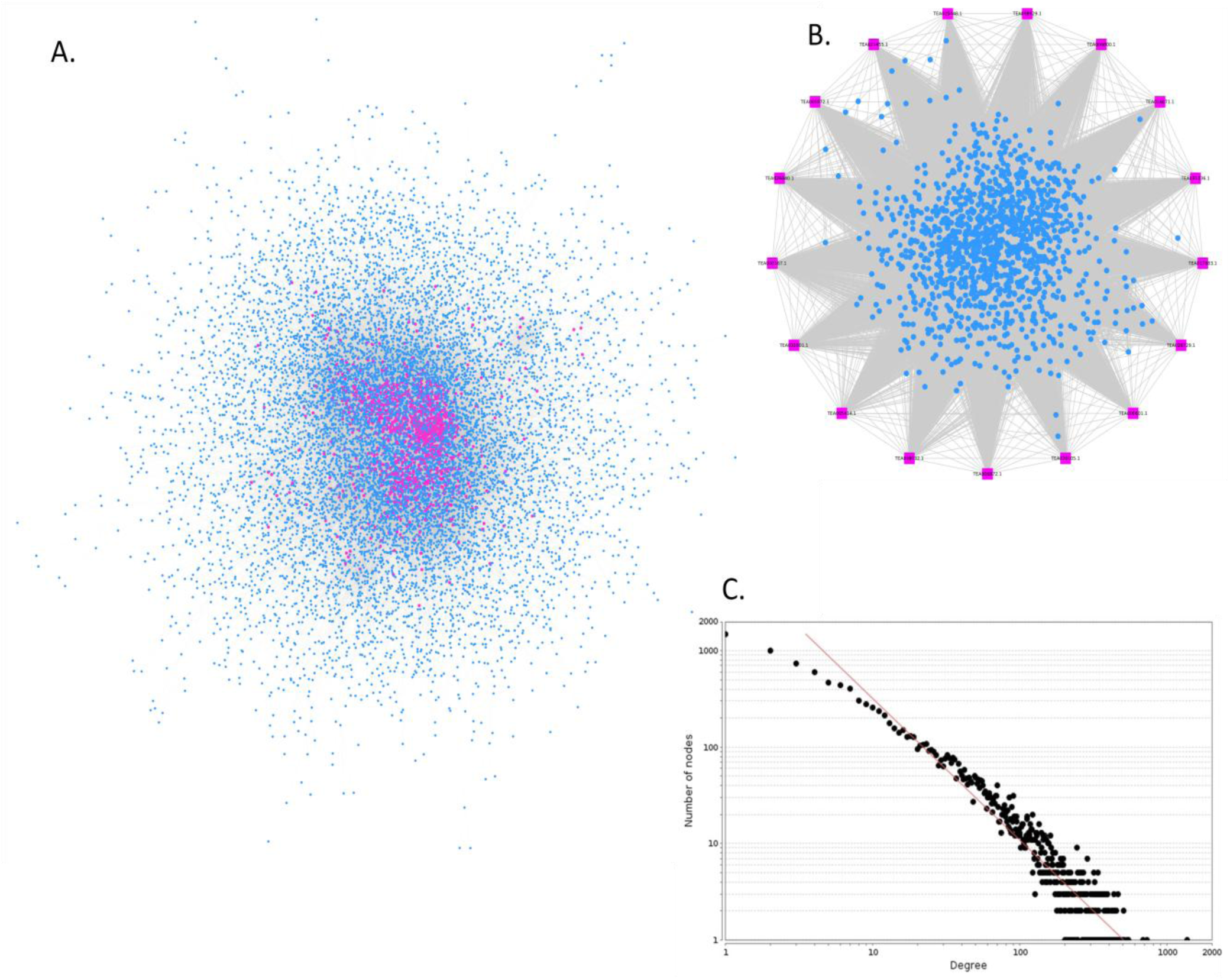
A. The genome wide interologous network of tea (TeaGPIN) with all key proteins (in purple). B. Highly influential key proteins (purple; rectangular) predicted using three centrality measures against both the sets of random networks. C. Degree distribution of the TeaGPIN with power-law fit (exponent is −1.47).

#### 3.1.1. Domain interaction based prioritization of tea interologs

Proteins interact through specific domains. Therefore, to assess the domain-domain interactions in all the interologs of TeaGPIN, participating proteins were examined for the presence of various domains. A total of 8,398 proteins were predicted to have 2,503 unique domains. These domains were further mapped to domain-domain interaction (DDI) data extracted from DOMINE database and 22,344 domain-domain interactions were successfully predicted among the interologs of TeaGPIN. All the interactions predicted in TeaGPIN were ranked using a confidence score S_total_. Interactions with S_ilog_ < 3 and S_dip_ = 0, were considered as low confidence interologs (LCI) and were filtered out as they were obtained from a small number of template plants and did not have support by the DDI score.

A total of 121,333 interactions were found to fall in the LCI category and were removed. Further, the S_total_ score of remaining TeaGPIN interologs were normalized using min-max scaling that resulted in the normalized score having the range [0,1]. PPIs scoring between 0 and 0.1 (*i.e.* the lowest 10% of normalized S_total_) were categorized as moderate confidence interologs (MCI). We obtained 35,907 interologs having normalized S_total_ score less than 0.1. All the interactions scoring ≥ 0.1 of normalized S_total_ were called high confidence interologs (HCI) and the resulting network of these interologs was termed as high-confidence TeaGPIN (hc-TeaGPIN). A total of 58,867 interacting pair among 5,983 proteins were found to have normalized S_total_ ≥ 0.1 and its largest component consisting of 5,770 nodes and 58,711 interactions was further used to evaluate function based reliability of TeaGPIN (Supplementary table S2).

#### 3.1.2. Function based reliability assessment of hc-TeaGPIN

Since interacting proteins are most often found to be localized in the same sub-cellular location, additional confidence on each interaction in hc-TeaGPIN was obtained by examining the sub-cellular locations of both the proteins involved in each interolog. All the interologs of hc-TeaGPIN were assessed for their localization in the same sub-cellular compartment. Figure 3 depicts the distribution of number of PPIs in different sub-cellular locations for hc-TeaGPIN (dots with empty centres) alongside distribution of randomly generated PPI pairs. Each number in random distribution is an average over ensemble of 10,000 *G*_*n,m*_ type ER networks and bars represent their ranges. Results reveal a complete contrast between the two distributions *i.e.* between the sub-cellular localization distribution of hc-TeaGPIN interologs and that of average random ensemble. The number of PPI pairs with same locations in hc-TeaGPIN is found to be significantly higher for most of the locations as compared to their random counterparts.

**Figure 3.**
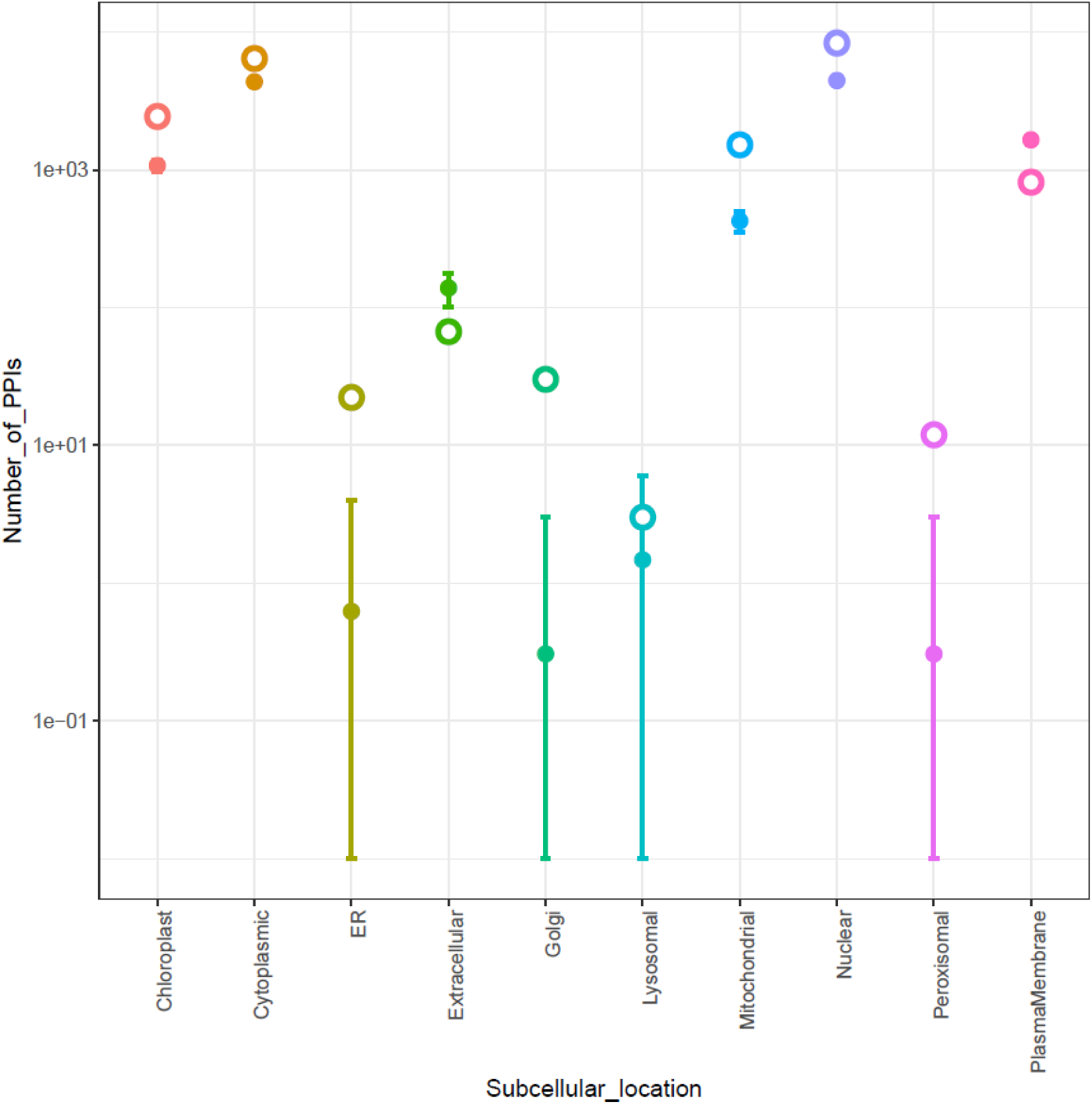
Protein co-localization similarity analysis of hc-TeaGPIN in 10 sub-cellular compartments. PPI frequencies of hc-TeaGPIN are represented by open circles and the frequencies of PPIs in 10,000 realizations of ER type random networks corresponding to hc-TeaGPIN are represented by filled circles, bars representing the ranges of frequencies.

### 3.2. Identification of key proteins in TeaGPIN

For the identification of proteins involved in key structural properties of TeaGPIN, we selected the giant component of network (11,916 nodes and 216,037 interactions) leaving the other nodes that were present in small independent clusters. This giant component of the TeaGPIN was used to construct two sets of random networks having similar dimensions; first set consists of 10,000 realizations of *G*_*n,m*_ type Erdős-Rényi model and the second set consists of 10,000 realizations of scale free models based on an extended Barabási-Albert algorithm. Based on three centrality measures, namely, betweenness, degree and eigenvector, a total of 1,302 key proteins having 56,724 interactions were identified having *p-*value ≤ 0.01 (Figure 2B). Among key proteins, 1,072 proteins have important regulatory role in 308 pathways. All the key proteins with their respective KO ids and the number of interactions they are involved-in are shown in Supplementary table S3. Giant component of the TeaGIPIN with key regulating proteins is shown in Figure 2A.

Protein TEA021735.1 has 1,361 interactions found in Module_1 that has important role in protein kinase activity and might be involved in regulatory processes in reproduction through glucose signalling [41]. Protein TEA006404.1 having KO id K13998 was found to have 728 interactions, and may have an important role in nucleotide metabolism in response to drought and heat stresses [42]. While protein TEA011031.1 having KO id K12860 was found to have 686 interactions and lies in spliceosome pathway under genetic information processing. Spliceosome is a complex of ribonucleoproteins with crucial roles in intron removal or splicing of mRNAs to regulate the transcription [43].

#### 3.2.1. Identification of highly influential key proteins

To obtain the set of key proteins, reported in the abovementioned section, two sets of important proteins corresponding to the ER and SF random networks were developed having *p-*value ≤ 0.01 in either of the three network metrics. Proteins that fall in all the six subsets, are termed as highly influential key proteins. For that, all the proteins which are common to these six subsets were further analysed as these may be the novel candidates for many processes. A total of 17 proteins were predicted having 1,223 interactions with highest degree of 540 of protein TEA008032.1 followed by TEA021455.1 and TEA000167.1 proteins with degrees 504 and 494, respectively. Protein TEA008032.1 encodes for ribosomal protein L3 (RPL3) which is related to plant growth and development by regulating translation and ribosome biogenesis [44]. Protein TEA021455.1 is predicted to be a U3 small nucleolar ribonucleoprotein protein (IMP3), these types of proteins are associated with the pre-ribosomal RNA processing and preribosomes assembly [45]. While protein TEA000167.1 has an important function in the encoding of 60S ribosomal protein L4 like, this protein has important role during cold stress as reported earlier in *Arabidopsis* and might be crucial for cold acclimation in tea as well [46]. All these proteins were also found to be interacting with two proteins related to flavonoid pathway (TEA027386.1 and TEA009431.1) that might be novel proteins and can be selected for the studies involving the extraction of important tea compounds. All of these highly influential key proteins along with their first degree interactors are shown in Figure 2B.

### 3.3. Gene ontology annotation and pathway analysis of TeaGPIN proteins

Through gene ontology annotation 12,194 GO terms were successfully associated with 6,341 node proteins of TeaGPIN. A total of 3,873 (∼31%) GO terms were classified into 34 categories of “biological processes”. Further, 20 categories were identified as “cellular components” with 3,604 (∼29%) GO terms and 34 categories from “molecular functions” were identified with 4,717 (∼38%) GO terms. Among the class of “biological processes”, ‘cellular process and metabolic process’ category was found to be most enriched followed by ‘response to stimulus’ and ‘biological regulation’. ‘Cell and cell part’ was highly enriched followed by ‘membrane’ and ‘protein-containing complex’ in “cellular components” class. Within “molecular functions” class, ‘catalytic activities and transporter activity’ was the most enriched category followed by ‘antioxidant activity’ and ‘structural molecular activity’ (Figure 4). Gene ontology (GO) term assignment to a large number of genes involved in TeaGPIN indicates towards the existence of diversified gene families in *Camellia sinensis.* Moreover, for finding the crucial regulatory pathways, a total of 392 pathways were identified by assigning 2,918 unique KO ids to 5,184 transcripts by means of KEGG database. Pathways obtained from KEGG database were further divided into five major categories which involves Metabolism, Genetic information processing, Environmental information processing, Cellular processes and Organismal systems as shown in Figure 5. Metabolism category is further categorized into 11 metabolic pathways with 535 unique transcripts in carbohydrate metabolism followed by amino acid metabolism (358 transcripts). For growth and development of plant systems, enzymes have important role in regulation of pathways by initiating specific mechanism related to growth and development of plants at particular stress condition [47]. Carbohydrate metabolism and amino acid metabolism are directly associated with changes in primary metabolism of plants that are mainly caused due to the biotic stresses. These act as precursors for the initiation and regulation of plant metabolites related to defence in order to counter stress conditions [48].

**Figure 4.**
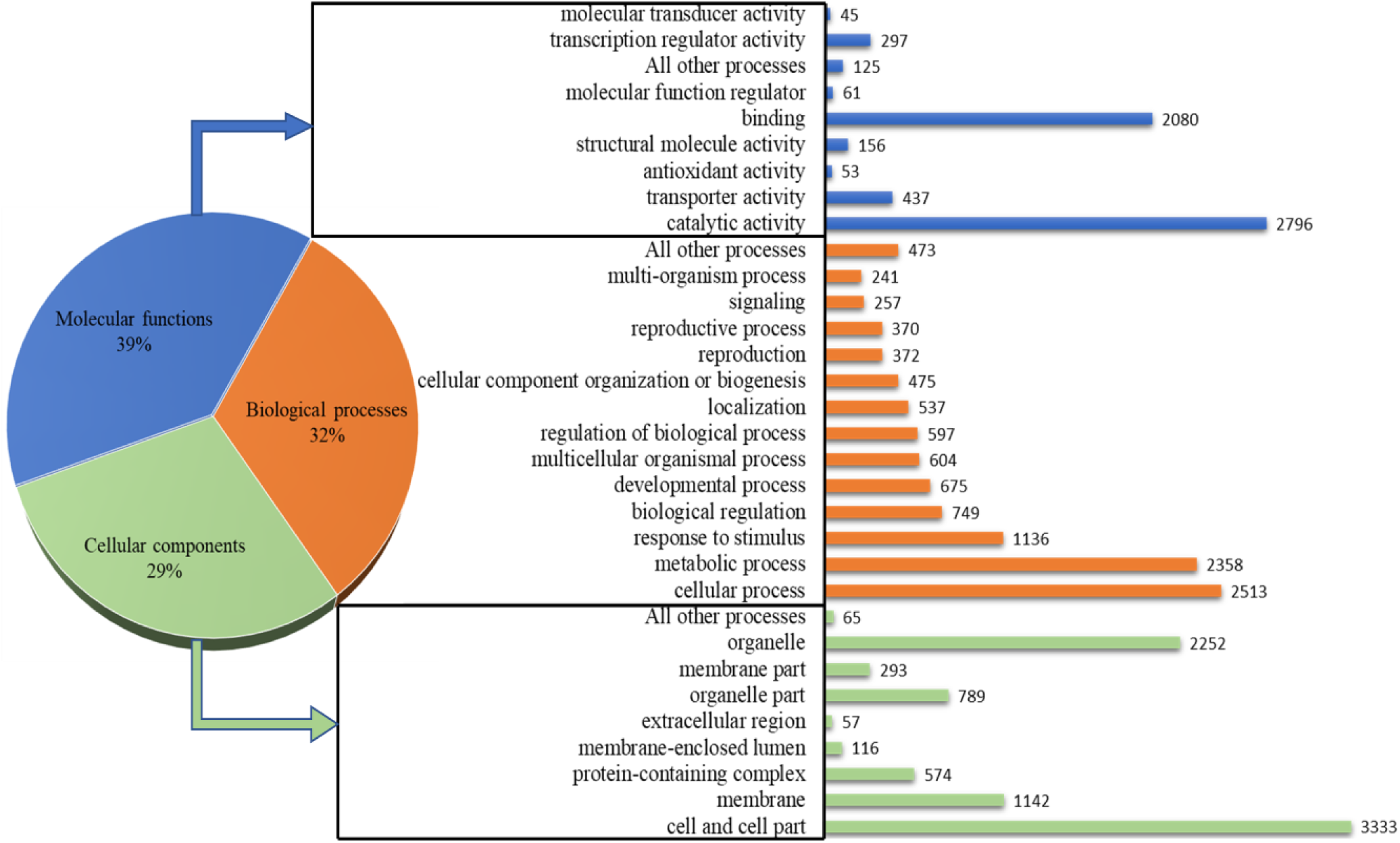
Gene Ontology (GO) annotation based characterization of proteins participating in TeaGPIN.

**Figure 5.**
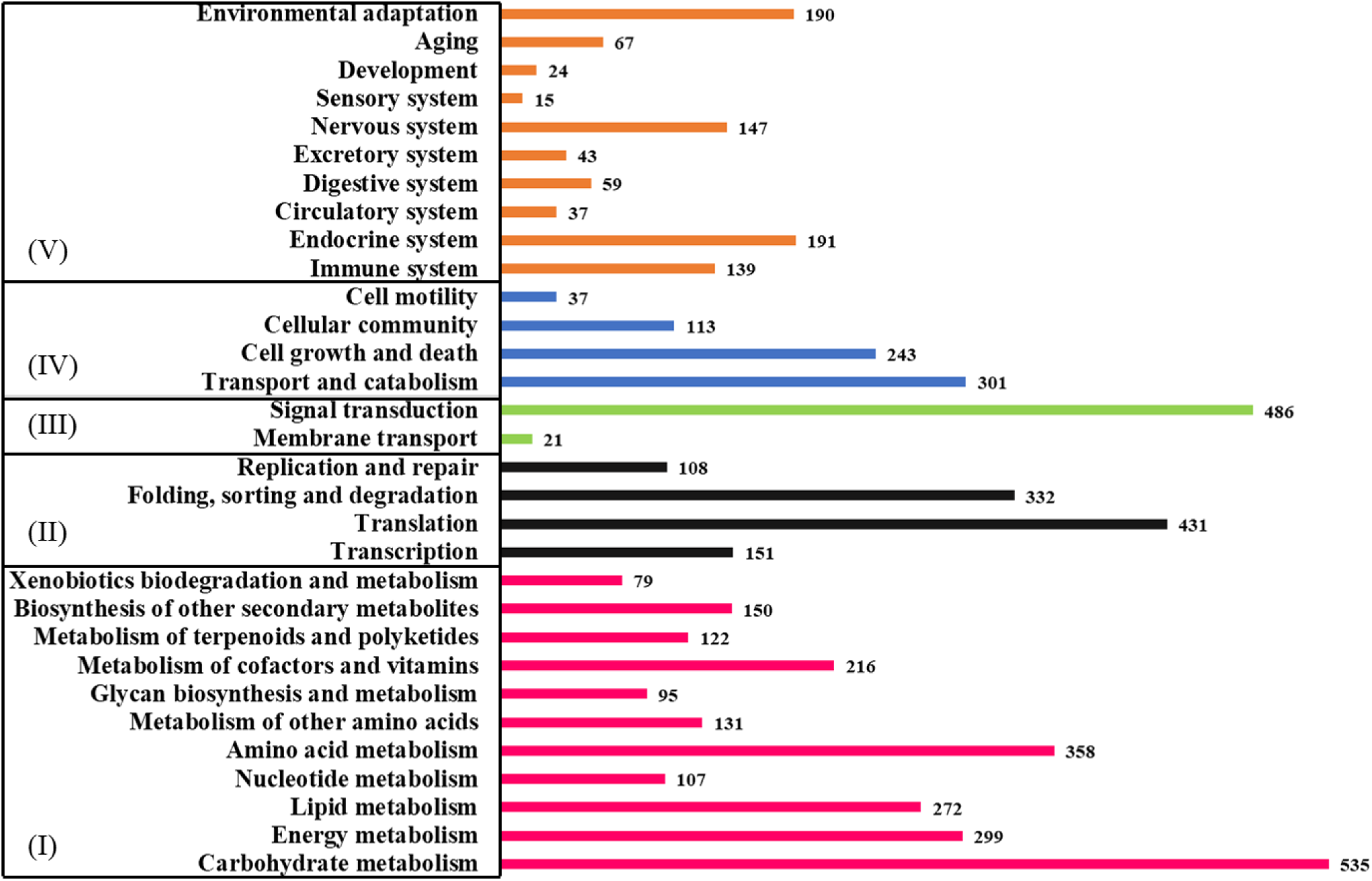
Pathway categorization of TeaGPIN proteins. (I) Metabolism, (II) Genetic information processing, (III) Environmental information processing, (IV) Cellular processes and (V) Organismal systems.

In another category of genetic information processing, a total of 431 genes were found in translation followed by 332 genes in folding, sorting and degradation, 151 genes in transcription and 108 in replication and repair. To identify the specific genes involved in hormonal signaling, pathways related to translational signaling at molecular level must be activated [49]. On the other hand several types of biotic and abiotic stresses can interfere in the functioning of proteins which leads to change in folding and accumulation of unfolded proteins, under this situation folding, sorting and degradation represents those heat shock proteins for the integrity of functional mechanism [50]. For particular conditions through transcription regulation, signal transduction pathways mechanism has crucial role in triggering the translational regulations by intricate feedback regulations [49]. Replication and repair have important role in maintaining the integrity of the genome through several checkpoints of different regulators at stress conditions during growth and development of plant [51]. In environmental information processing, two processes ‘signal transduction’ and ‘membrane transport’ were found with 486 and 21 genes, respectively. To adapt the stressful conditions under continuously fluctuating environment, signal transduction has critical role as it assesses the stress condition and helps in adapting to the particular one [52]. Membrane transport systems are very important in maintaining the homeostasis of cell during ion-transport activity which is responsible for stress condition [53]. Four processes, *namely*, transport and catabolism, cell growth and death, cellular community and cell motility were found to have 301, 243, 113 and 37 genes, respectively. Transport and catabolism has important role in defence by recognizing the specific receptors of binding ligands from the infectious agents that helps in remodeling the tissue during any pathogen attack [54]. In response to stress from the external environmental factors, cell death and cell motility is the process to have an important role in the development of cell to survive by sensing chemical gradients [55]. In the organismal systems category, the functions of plant systematics can be predicted, a total of 10 processes were successfully identified with highest number of genes (191) in endocrine system followed by environmental adaptation (190). Other processes like, immune systems and aging were also predicted that may be having vital roles during stress conditions for survival of the plant.

### 3.4. Identification of transcription factors (TFs) in TeaGPIN

Transcription factors have very important and specific role in various signalling and cellular processes in response to any stress condition during growth. We attempt to screen transcription factors from all 12,033 interacting proteins of TeaGPIN and report a total of 540 transcription factors by using two methods (Figure 6). Firstly, by downloading all sequences belonging to transcription factors from plant transcription factor database, we have developed 58 HMMs by developing an alignment of all the sequences of a particular category, and secondly, by using iTAK tool. We have selected those sequences as transcription factors which are common in both the methods. Among 540 predicted transcription factors, 59 are related to MYB or MYB_related followed by C2H2 (39), GRAS (35) and bHLH (32) and several other TF families. Various complexes of MYB/bHLH transcription factors are responsible for regulation of many processes that include cell death, cell wall synthesis or synthesis of specific metabolites, circadian clock, hormonal signalling, biotic and abiotic stress responses. In other words, interactions between bHLH and MYBs are based on the demand of cell at particular condition to regulate the cellular function [56–58]. Other transcription factors such as WRKY (16 TFs) are responsible for regulation of cell during biotic stress specifically during pathogen attack and also reported in other studies specific to tea [59]. NAC (27 TFs) are helpful in managing ROS load during stress conditions (biotic and abiotic stress), and may be crucial for tea as these have also been reported to have important regulatory roles in rice [60].

**Figure 6.**
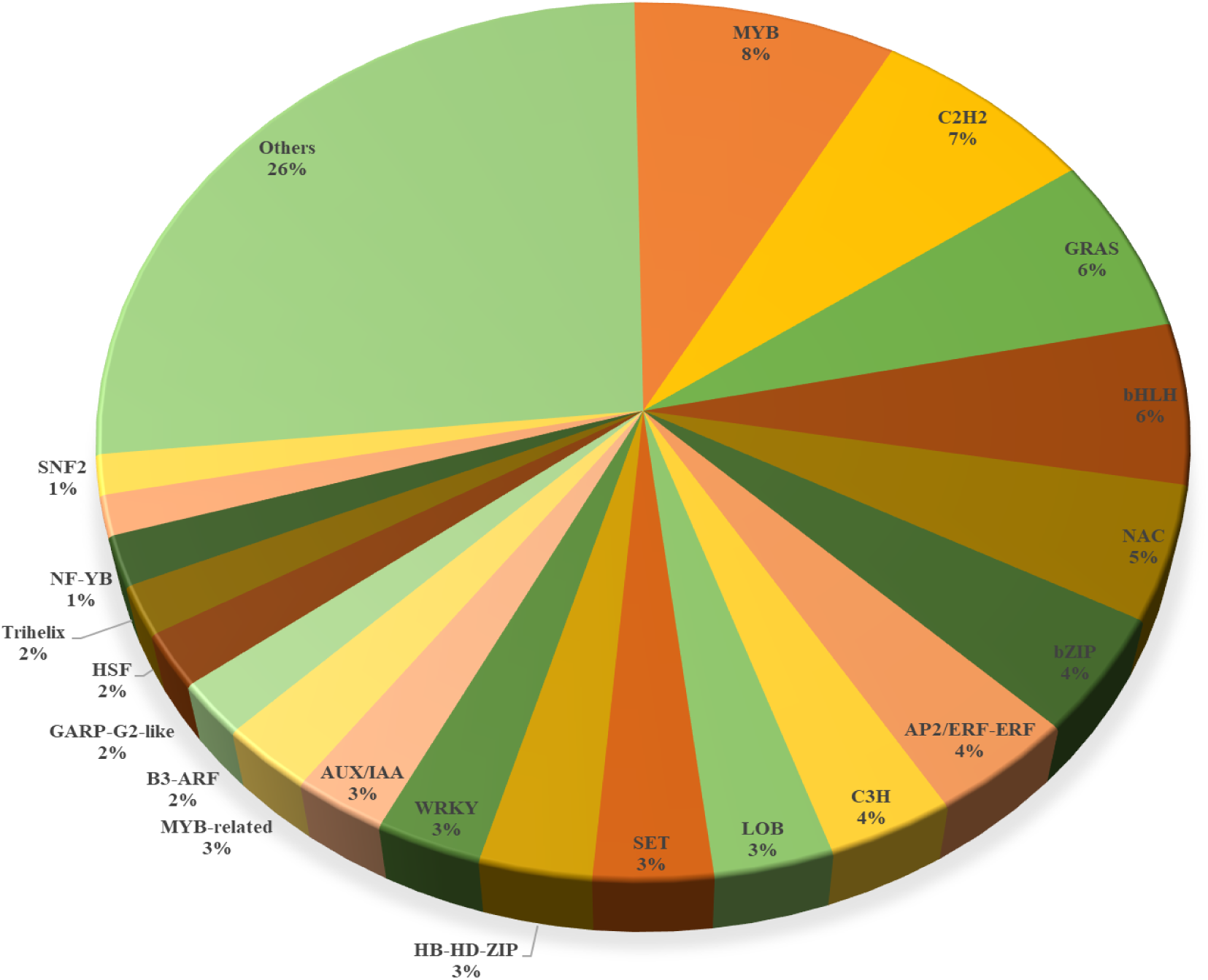
Classification of transcription factors characterized in the TeaGPIN proteins.

#### 3.4.1. Transcription factors in the key proteins of TeaGPIN

Among the key proteins, 8 proteins are found to be associated with transcription factor activities and have 505 interactions. Two proteins, namely, TEA025288.1 and TEA015567.1 are predicted to be the transcription factors of SNF2 family with KO id K19001 and K20098, respectively. TFs of SNF2-family have the ability of reducing the replication stress-induced DNA breaks and chromosomal aberrations as well as restoring the replication fork stability by remodelling stalled forks [61]. ProteinTEA031026.1 encodes for a C3H transcription factor with KO id K14404; C3H transcription factors are reported to have regulatory roles in plants in response to stimulation by multiple signals [62]. Two other TF families having key proteins of TeaGPIN are MYB_related (TEA021255.1; 14 interactions) and LFY (TEA003201.1; 19 interactions). MYB_related transcription factors have crucial role to several stresses by hormonal stimuli [63], while LFY transcription factors are known to act as central regulators of crucial developmental processes in plants that include reproduction and floral transition [64].

#### 3.4.2. TeaGPIN based characterization of SNF2 and C3H transcription factor families

As a case study, we have selected the SNF2 and C3H transcription factor families as their detailed characterization is still lacking in tea. In this section, we attempt a network based exploration of the proteins involved in these two families and identify their potential implications in important regulatory processes (Figure 7). SNF2 family TFs have roles in transcription regulation and are also important for maintaining chromosome stability during mitosis. These are known to be crucial during various processes of DNA damaging such as post-replication daughter strand gap repair, nucleotide excision repair and recombinational pathways [65]. A total of 8 proteins of SNF2 family were mapped in TeaGPIN and were having 289 interactions. Among these, protein TEA025288.1 was found to have the highest degree of 105 followed by TEA000359.1 having 103 interactions. Two proteins of this family, namely, TEA000359.1 and TEA025288.1 were found to be in the key proteins having 33 and 39 interactions, respectively. During pathway analysis of these key proteins (TEA000359.1; K10841 and TEA025288.1; K19001), it is found that these are involved in folding, sorting and degradation pathway with important roles in regulating nucleotide excision repair mechanism and DNA repair and recombination.

**Figure 7.**
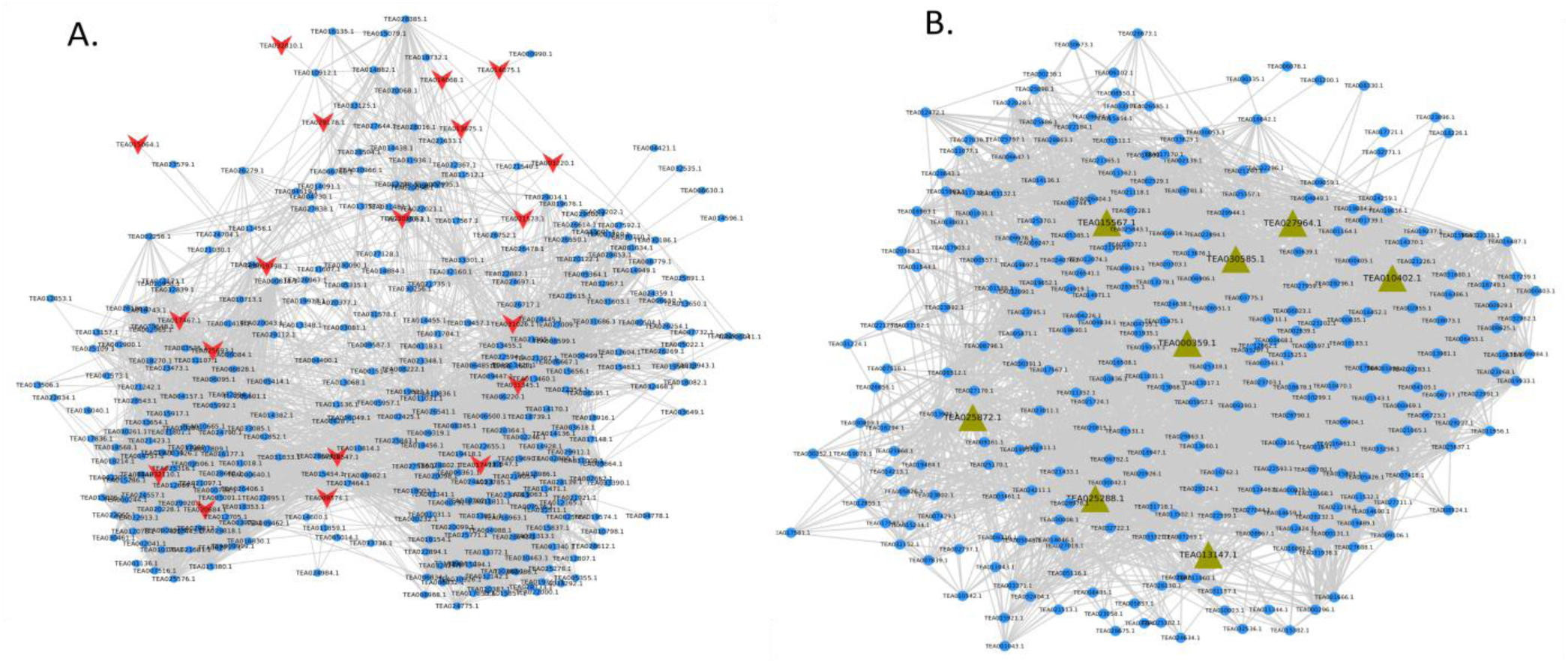
Subnetworks of A. transcription factors belonging to C3H family (purple; V shaped), and B. transcription factors belonging to SNF2 family (green; triangular shaped)

Proteins belonging to C3H family contain zinc finger associated with WD40 repeats at the end terminal part of the protein. C3H zinc finger type of motif occurs in several proteins ranging from yeast to human. Due to their transcriptional activity, these TFs regulate RNA binding proteins to induce RNA processing in response to drought stress [66]. A total of 19 proteins having 351 interactions were determined with highest degree 118 of protein TEA031026.1 followed by protein TEA025693.1 with degree 78 and protein TEA031345.1 with degree 64. Furthermore, protein TEA031026.1 was also found to be in the list of key proteins. During pathway analysis, it has been found that all these proteins have roles in genetic information and processing pathways to regulate important mechanism related to growth and development of plant.

### 3.5. Modular architecture of TeaGPIN

Several complex biological processes are continuously occurring within the cell through various interactions among specific proteins, thus, to discover the knowledge about underlying subcellular mechanisms for specific biological processes, identification and cataloging of functional modules in the systems scale PPI network is an important step. MCODE assisted clustering analysis was performed for TeaGPIN, and a total of 206 modules consisting of 4,150 proteins were obtained [Supplementary table S4]. Furthermore, on the basis of maximum number of nodes in each cluster, we have selected 10 modules (1,658 nodes and 31,464 interactions) for further analysis.

Module_1 includes 223 proteins in which metabolic pathways are found to be highly enriched followed by biosynthesis of secondary metabolites and biosynthesis of antibiotics. About 92 proteins with total of 1,805 interactions are found in metabolic pathways, 58 proteins with 1,318 interactions in biosynthesis of secondary metabolites and 34 proteins with 933 interactions in biosynthesis of antibiotics. Protein TEA033193.1 (161 interactions) followed by protein TEA000324.1 (149 interactions) and protein TEA020428.1 (129 interactions) have highest interactions in all the pathways of Module_1.

Module_2 consists of 195 proteins, among these 29 proteins with 674 interactions are involved in enriching aminoacyl-tRNA biosynthesis pathway in which proteins TEA028953.1, TEA017370.1 and TEA021820.1 have 196, 166, and 161 interactions, respectively. A total of 17 proteins with 510 interactions were found to be enriching the photosynthesis pathway in which proteins TEA018071.1, TEA022973.1 and TEA031434.1 have 284, 265 and 256 interactions, respectively. A total of 15 proteins with 528 interactions have role in DNA replication pathway enrichment with proteins TEA002561.1, TEA010299.1 and TEA031525.1 having318, 234 and 228 interactions, respectively.

In the Module_3, total 191 proteins are found enriching ribosomal biogenesis, ribosomes and RNA transport pathway. In ribosomal biogenesis, 29 proteins with 963 interactions are found in which proteins TEA026406.1 and TEA000640.1 both have 433 interactions followed by the protein TEA029818.1 having 391 interactions. In ribosomes pathway, 23 proteins having 745 interactions are found, in which proteins TEA012427.1, TEA006558.1 and TEA017734.1 have 368, 250 and 217 interactions, respectively. RNA transport pathway in Module_3 consists of 15 proteins with 766 interactions, where proteins TEA021097.1, TEA021242.1 and TEA025964.1 have 306, 257 and 252 interactions, respectively.

Additionally, KEGG pathway enrichment of all the proteins involved in 10 most populated modules was performed and it resulted in the identification of 1,261 proteins with 920 unique KO ids. In translation category, under genetic information processing, ribosome pathway is highly enriched with 140 proteins (20 proteins from Module_3, 119 proteins from Module_4 and 1 protein from Module_10), RNA transport pathway is found to be highly enriched with 46 proteins (8 proteins from Module_1, 2 proteins from Module_2, 14 proteins from Module_3, 12 proteins from Module_4, 3 proteins from Module_7, 5 proteins from Module_10, 1 protein each from Module_6 and Module_8) in translation. In the process of ribosome biogenesis, various ribosome biogenesis factors (RBFs) are involved in mainly three components of the cell (nucleolus, nucleus and cytoplasm) for primary ribosomal processing [67]. Various cellular processes are directly controlled by RNA transport as it has wide role from embryo development to activation of defence system in plants by several signaling cascades [68]. Also, in metabolism, oxidative phosphorylation category under energy metabolism is enriched with 44 proteins (17 proteins from Module_9, 10 proteins from Module_5, 5 proteins from Module_7, 4 proteins each from Module_2 and Module_6, 1 protein from Module_3 and Module_1 and 2 proteins from Module_10). Oxidative phosphorylation has important role in regulating cellular processes like respiration and ATP biosynthesis for plant metabolic activities [69].

### 3.6. Sub-networks of proteins involved in flavonoid biosynthesis and photosynthesis pathways

In the secondary metabolites of tea, flavonoids are considered as key metabolites having significant contribution to enhance the industrial or agronomic value of plant product [70]. Specifically, in the case of green tea, flavonoids are the major antioxidative constituents of tea leaves having various health benefits [71]. We attempt to identify the proteins involved in flavonoid biosynthetic pathway using KEGG database and available literature. Total 31 proteins could be identified that were having 399 interactions in the TeaGPIN, among these, 12 are identified as key proteins interacting with 887 proteins.

Three proteins, namely, TEA009431.1, TEA006577.1 and TEA034012.1 have 4-coumarate CoA ligase activity (4CL) with KO id K01904 and are found to have 228, 71 and 63 interactions, respectively. 4CL is synthesized due to catalytic activity of p-coumarate: CoA ligase by using the substrates as acetyl-CoA and 4-coumaric acid, 4CL has important role in catalysing the ligation of CoA to hydroxycinnamic acids which is a crucial point in order to regulate monolignol or flavonoid pathways [72,73]. Protein TEA014056.1 is found to have 65 interactions and has KO id K10775 encoding for phenylalanine ammonia-lyase (PAL). PAL is an important enzyme for the synthesis of Phenylpropanoids which engenders various types of aromatic metabolites that are crucial for environmental adaptation, development and growth of plant [74]. PAL is also important for salicylic acid biosynthesis by pathogen-induced trigging as reported in soybean and might also be important for initiating defence mechanism in tea in response to biotic stress [75].

Protein TEA023790.1 with 55 interactions and KO id K00475 codes for F3H enzyme (Flavanone 3-hydroxylase). F3H is a known nuclear enzyme important for flavonoid biosynthetic pathway by catalysis of 3-hydroxylation of (2S)-flavanones that includes naringenin to dihydroflavonols [76]. Protein TEA032730.1 with 50 interactions has function of bifunctional dihydroflavonol 4-reductase (DFR) that is an oxidoreductase and produces flavan 3,4-diols by catalysing the NADPH dependent reduction of keto group at the 4^th^ position of dihydroflavonols. Flavan 3,4-diols are the precursors to form anthocyanidins and flavan 3-ols that form condensed tannins. DFR is important in the formation of three classes of flavonoids which includes anthocyanin pigments, flavanols responsible for abiotic stresses and flavanols which act as sunscreen [77]. Proteins involved in flavanol synthesis are shown in Figure 8A.

**Figure 8.**
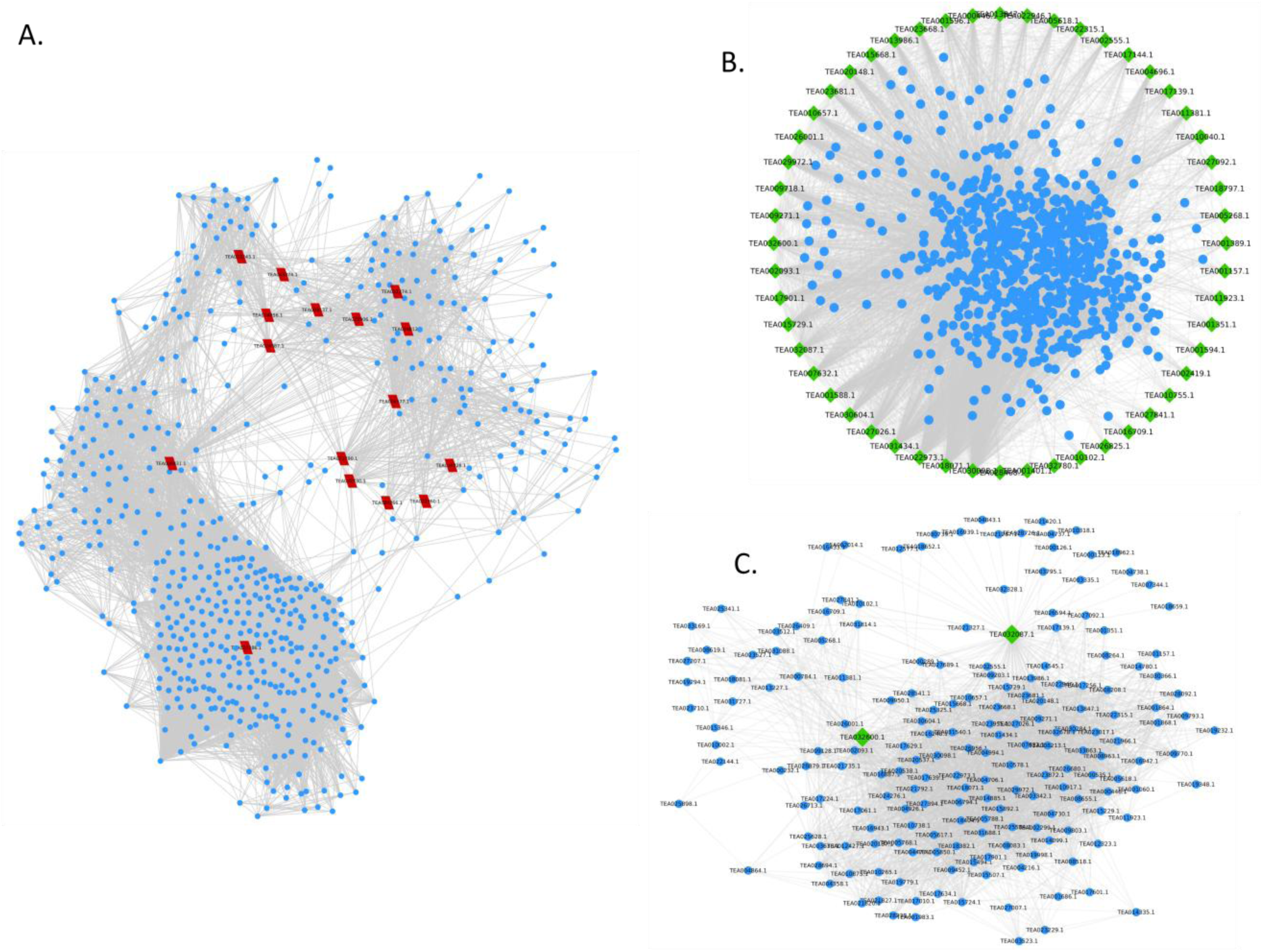
Subnetworks of proteins involved in A. flavonoid biosynthesis pathway (olive; parallelogram shaped) and B. photosynthesis pathway (light green colour; dimond shaped). C. Protein interactors of two important proteins of photosynthesis, namely, plastocyanin (Pc) and Ferredoxin NADP+ reductase (FNR), both highlighted in green..

Abiotic stress has direct impact on photosynthesis, as all plants (depending upon their types, C3 or C4) respond to the stress conditions with various alterations of gene expressions and other environmental factors [78]. TeaGPIN has been analyzed to unravel key proteins involved in photosynthetic process (Figure 8B). A total of 52 proteins were found to be interacting with 770 proteins with highest number of interactions in protein TEA030098.1 (340 interactions) encoding ATPF1D (F-type H+ transporting ATPase subunit delta) and protein TEA018071.1 (284 interactions) encoding ATPF1G (F-type H+ transporting ATPase subunit gamma). F-type ATPases have important role in catalysing the synthesis of ATP through coupled rotary motors complexes F_0_ and F_1_ [79]. Further, other proteins examined in the photosynthetic pathway are TEA032087.1 (plastocyanin; Pc) and TEA032600.1 (Ferredoxin NADP+ reductase; FNR) with 124 and 102 interactions, respectively in TeaGPIN (Figure 8C). Pc is a soluble protein and is located in the thylakoid lumen of chloroplast, its main role is to mediate photosynthetic electron transport among two complexes (cytochrome b_6_f and photosystem I) of membrane proteins [80]. Plastocyanin is earlier reported to have indispensable role of mediating electron flow during photosynthesis in *Arabidopsis thaliana* [81]. FNR is the member of NADPH dependent oxidoreductases of the flavoprotein superfamily; its important role is to participate in transfer of electrons from reduced ferredoxin to the NADPH, and in the process providing a reductant for carbon fixation in plants [82].

## Summary

Tea is very important crop due to its rich taste and medicinal values and has high commercial value. Continuously changing environmental conditions and various pathogens are threat for the production and quality of this crop. Thus, to identify the proteins involved in important pathways and for their potential implications in delineating their mechanisms of actions, we have developed genome wide interolog based protein-protein interaction network (TeaGPIN) with 12,033 nodes and 216,107 interactions. Furthermore, a high confidence TeaGPIN (hc-TeaGPIN) consisting of 5,983 nodes and 58,867 interactions is constructed from the complete network by prioritizing the interactions using a hybrid approach involving domain-domain interaction score and interolog score. Further, the biological significance of each high confidence interaction was assessed by computing protein co-localization similarity (PCS). PCS values were found high for most of the interologs as compared to ER type random networks. Furthermore, by comparing the complete TeaGPIN with 10,000 random realizations of ER (*G*_*n,m*_ type) and 10,000 of SF (extended BA type) models, a total of 1,302 key proteins were identified which may be involved in crucial processes of tea growth, development and pathology. Additionally, TeaGPIN is sought to identify small independent functional units called modules and most of the modules are found to be involved in secondary metabolites and antibiotics biosynthesis that is further confirmed by GO annotations.

We have also conducted three case studies towards network based characterization of the transcription factor families and proteins involved in flavonoid biosynthesis and photosynthesis pathways. In the future, this network (TeaGPIN) can be explored to identify various proteins and their interactors related to growth, development and response to immune system. This methodology can also be implemented for newly sequenced genomes of other plants to understand the complexity of systems.

## Authors Contributions

VS* conceptualized and supervised the study. GS and VS^†^ performed computational experiments. All the authors analyzed and interpreted the results as well as wrote the manuscript.

## Conflict of interest

All the authors declare that there is no conflict of interest.

## Acknowledgement

All the authors thank Central University of Himachal Pradesh for providing the required facilities. VS^†^ thanks Council of Scientific and Industrial Research (CSIR), India for the junior research fellowship (JRF).

## References

[1] P. Braun, A. Gingras, History of protein–protein interactions: From egg-white to complex networks, Proteomics. 12 (2012) 1478–1498.

[2] L. Cuadra, S. Salcedo-Sanz, J. Del Ser, S. Jiménez-Fernández, Z. Geem, A critical review of robustness in power grids using complex networks concepts, Energies. 8 (2015) 9211–9265.

[3] P. Braun, S. Aubourg, J. Van Leene, G. De Jaeger, C. Lurin, Plant Protein Interactomes, Annu. Rev. Plant Biol. 64 (2013) 161–187.

[4] A.-L. Barabasi, Z.N. Oltvai, Network biology: understanding the cell’s functional organization, Nat Rev Genet. 5 (2004) 101–113.

[5] R. Thanasomboon, S. Kalapanulak, S. Netrphan, T. Saithong, Prediction of cassava protein interactome based on interolog method, Sci. Rep. 7.1 (2017) 17206.

[6] D. Di Silvestre, A. Bergamaschi, E. Bellini, P. Mauri, Large Scale Proteomic Data and Network-Based Systems Biology Approaches to Explore the Plant World, Proteomes. 6 (2018) 27.

[7] R. Sharan, I. Ulitsky, R. Shamir, Network-based prediction of protein function, Mol. Syst. Biol. 3 (2007) 88.

[8] U. Stelzl, U. Worm, M. Lalowski, C. Haenig, F.H. Brembeck, H. Goehler, M. Stroedicke, M. Zenkner, A. Schoenherr, S. Koeppen, A human protein-protein interaction network: a resource for annotating the proteome, Cell. 122 (2005) 957–968.

[9] K. Tarassov, V. Messier, C.R. Landry, S. Radinovic, M.M.S. Molina, I. Shames, Y. Malitskaya, J. Vogel, H. Bussey, S.W. Michnick, An in vivo map of the yeast protein interactome, Science 320 (2008) 1465–1470.

[10] P. Uetz, L. Giot, G. Cagney, T.A. Mansfield, R.S. Judson, J.R. Knight, D. Lockshon, V. Narayan, M. Srinivasan, P. Pochart, A comprehensive analysis of protein–protein interactions in Saccharomyces cerevisiae, Nature. 403 (2000) 623.

[11] E. Formstecher, S. Aresta, V. Collura, A. Hamburger, A. Meil, A. Trehin, C. Reverdy, V. Betin, S. Maire, C. Brun, Protein interaction mapping: a Drosophila case study, Genome Res. 15 (2005) 376–384.

[12] J. Geisler-Lee, N. O’Toole, R. Ammar, N.J. Provart, A.H. Millar, M. Geisler, A Predicted Interactome for Arabidopsis, PLANT Physiol. 145.2 (2007) 317–329.

[13] H. Gu, P. Zhu, Y. Jiao, Y. Meng, M. Chen, PRIN: A predicted rice interactome network, BMC Bioinformatics. 12.1 (2011) 161.

[14] G. Zhu, A. Wu, X.-J. Xu, P.-P. Xiao, L. Lu, J. Liu, Y. Cao, L. Chen, J. Wu, X.-M. Zhao, PPIM:A Protein-Protein Interaction Database for Maize, Plant Physiol. 170.2 (2016) 618–626.

[15] B. Alipoor, A.H. Rad, A review on the therapeutical effects of tea, Asian J. Clin. Nutr. 4 (2012) 1–15.

[16] G. Singh, V. Singh, V. Singh, Construction and analysis of an interologous protein–protein interaction network of Camellia sinensis leaf (TeaLIPIN) from RNA–Seq data sets, Plant Cell Rep. (2019) 1–14. doi: 10.1007/s00299-019-02440-y

[17] W.-L. Wang, Y.-X. Wang, H. Li, Z.-W. Liu, X. Cui, J. Zhuang, Two MYB transcription factors (CsMYB2 and CsMYB26) are involved in flavonoid biosynthesis in tea plant [Camellia sinensis (L.) O. Kuntze], BMC Plant Biol. 18 (2018) 288.

[18] C. Wei, H. Yang, S. Wang, J. Zhao, C. Liu, L. Gao, E. Xia, Y. Lu, Y. Tai, G. She, Draft genome sequence of Camellia sinensis var. sinensis provides insights into the evolution of the tea genome and tea quality, Proc. Natl. Acad. Sci. 115 (2018) E4151–E4158.

[19] E.-H. Xia, H.-B. Zhang, J. Sheng, K. Li, Q.-J. Zhang, C. Kim, Y. Zhang, Y. Liu, T. Zhu, W. Li, The tea tree genome provides insights into tea flavor and independent evolution of caffeine biosynthesis, Mol. Plant. 10 (2017) 866–877.

[20] D. Szklarczyk, A.L. Gable, D. Lyon, A. Junge, S. Wyder, J. Huerta-Cepas, M. Simonovic, N.T. Doncheva, J.H. Morris, P. Bork, L.J. Jensen, C. Von Mering, STRING v11: Protein-protein association networks with increased coverage, supporting functional discovery in genome-wide experimental datasets, Nucleic Acids Res. 47 (2018) D607–D613.

[21] M.E. Smoot, K. Ono, J. Ruscheinski, P.-L. Wang, T. Ideker, Cytoscape 2.8: new features for data integration and network visualization, Bioinformatics. 27 (2010) 431–432.

[22] S. El-Gebali, J. Mistry, A. Bateman, S.R. Eddy, A. Luciani, S.C. Potter, M. Qureshi, L.J. Richardson, G.A. Salazar, A. Smart, E.L.L. Sonnhammer, L. Hirsh, L. Paladin, D. Piovesan, S.C.E. Tosatto, R.D. Finn, The Pfam protein families database in 2019, Nucleic Acids Res. 47 (2019) D427–D432.

[23] S. Yellaboina, A. Tasneem, D. V Zaykin, B. Raghavachari, R. Jothi, DOMINE: a comprehensive collection of known and predicted domain-domain interactions, Nucleic Acids Res. 39 (2010) D730–D735.

[24] V. Singh, G. Singh, V. Singh, TulsiPIN: an interologous protein interactome of Ocimum tenuiflorum, BioRxiv. (2019) 680025. doi:10.1101/680025.

[25] C.S. Yu, Y.C. Chen, C.H. Lu, J.K. Hwang, Prediction of protein subcellular localization, Proteins. 64 (2006) 643–651.

[26] H. Jeong, S.P. Mason, A.-L. Barabási, Z.N. Oltvai, Lethality and centrality in protein networks, Nature. 411 (2001) 41–42.

[27] M.P. Joy, A. Brock, D.E. Ingber, S. Huang, High-betweenness proteins in the yeast protein interaction network, Biomed Res. Int. 2005 (2005) 96–103.

[28] M.E.J. Newman, The mathematics of networks. The new palgrave encyclopedia of economics, (2008) 1–8.

[29] P. Erdös, A. Rényi, On random graphs, Publ. Math. Inst. Hung. Acad. Sci. 5.1 (1960) 17–61.

[30] A.-L. Barabási, R. Albert, Emergence of scaling in random networks, Science. 286 (1999) 509–512.

[31] N. Pržulj, D.G. Corneil, I. Jurisica, Modeling interactome: Scale-free or geometric?, Bioinformatics. 20 (2004) 3508–3515.

[32] T.Z. Berardini, L. Reiser, D. Li, Y. Mezheritsky, R. Muller, E. Strait, E. Huala, The Arabidopsis information resource: making and mining the “gold standard” annotated reference plant genome, Genesis. 53 (2015) 474–485.

[33] T. Tian, Y. Liu, H. Yan, Q. You, X. Yi, Z. Du, W. Xu, Z. Su, AgriGO v2.0: A GO analysis toolkit for the agricultural community, 2017 update, Nucleic Acids Res. 45 (2017) W122–W129.

[34] J. Ye, Y. Zhang, H. Cui, J. Liu, Y. Wu, Y. Cheng, H. Xu, X. Huang, S. Li, A. Zhou, X. Zhang, L. Bolund, Q. Chen, J. Wang, H. Yang, L. Fang, C. Shi, WEGO 2.0: A web tool for analyzing and plotting GO annotations, 2018 update, Nucleic Acids Res. 46 (2018) W71–W75.

[35] M. Kanehisa, M. Furumichi, M. Tanabe, Y. Sato, K. Morishima, KEGG: New perspectives on genomes, pathways, diseases and drugs, Nucleic Acids Res. 45 (2017) D353–D361.

[36] J. Jin, F. Tian, D.-C. Yang, Y.-Q. Meng, L. Kong, J. Luo, G. Gao, PlantTFDB 4.0: toward a central hub for transcription factors and regulatory interactions in plants, Nucleic Acids Res. 45 (2016) D1040–D1045.

[37] F. Sievers, A. Wilm, D. Dineen, T.J. Gibson, K. Karplus, W. Li, R. Lopez, H. McWilliam, M. Remmert, J. Söding, Fast, scalable generation of high-quality protein multiple sequence alignments using Clustal Omega, Mol. Syst. Biol. 7 (2011) 539.

[38] A. Krogh, B. Larsson, G. von Heijne, E.L. Sonnhammer, Predicting transmembrane protein topology with a hidden Markov model: application to complete genomes., J. Mol. Biol. 305 (2001) 567–580.

[39] Y. Zheng, C. Jiao, H. Sun, H.G. Rosli, M.A. Pombo, P. Zhang, M. Banf, X. Dai, G.B. Martin, J.J. Giovannoni, P.X. Zhao, S.Y. Rhee, Z. Fei, iTAK: A Program for Genome-wide Prediction and Classification of Plant Transcription Factors, Transcriptional Regulators, and Protein Kinases, Mol. Plant. 9.12 (2016) 1667–1670.

[40] G.D. Bader, C.W.V. Hogue, An automated method for finding molecular complexes in large protein interaction networks, BMC Bioinformatics. 4.1 (2003) 2.

[41] N. Govender, S. Senan, E.E. Sage, Z.-A. Mohamed-Hussein, M.M. Mackeen, R. Wickneswari, An integration of phenotypic and transcriptomic data analysis reveals yield-related hub genes in Jatropha curcas inflorescence, PLoS One. 13 (2018) e0203441.

[42] A. Das, P. Rushton, J. Rohila, Metabolomic profiling of soybeans (Glycine max L.) reveals the importance of sugar and nitrogen metabolism under drought and heat stress, Plants. 6 (2017) 21.

[43] L.J. Rivera-Vega, D.A. Galbraith, C.M. Grozinger, G.W. Felton, Host plant driven transcriptome plasticity in the salivary glands of the cabbage looper (Trichoplusia ni), PLoS One. 12 (2017) e0182636.

[44] S.C. Popescu, N.E. Tumer, Silencing of ribosomal protein L3 genes in N. tabacum reveals coordinate expression and significant alterations in plant growth, development and ribosome biogenesis, Plant J. 39 (2004) 29–44.

[45] D.J. Leary, M.P. Terns, S. Huang, Components of U3 snoRNA-containing complexes shuttle between nuclei and the cytoplasm and differentially localize in nucleoli: implications for assembly and function, Mol. Biol. Cell. 15 (2004) 281–293.

[46] Y. Kawamura, M. Uemura, Mass spectrometric approach for identifying putative plasma membrane proteins of Arabidopsis leaves associated with cold acclimation, Plant J. 36 (2003) 141–154.

[47] S. Mahajan, N. Tuteja, Cold, salinity and drought stresses: an overview, Arch. Biochem. Biophys. 444 (2005) 139–158.

[48] S. Zhou, Y.-R. Lou, V. Tzin, G. Jander, Alteration of plant primary metabolism in response to insect herbivory, Plant Physiol. 169 (2015) 1488–1498.

[49] C. Merchante, J. Brumos, J. Yun, Q. Hu, K.R. Spencer, P. Enríquez, B.M. Binder, S. Heber, A.N. Stepanova, J.M. Alonso, Gene-specific translation regulation mediated by the hormone-signaling molecule EIN2, Cell. 163 (2015) 684–697.

[50] C. Liu, R. Lu, G. Guo, T. He, Y. Li, H. Xu, R. Gao, Z. Chen, J. Huang, Transcriptome analysis reveals translational regulation in barley microspore-derived embryogenic callus under salt stress, Plant Cell Rep. 35 (2016) 1719–1728.

[51] Z. Hu, T. Cools, L. De Veylder, Mechanisms used by plants to cope with DNA damage, Annu. Rev. Plant Biol. 67 (2016) 439–462.

[52] C. Waszczak, S. Akter, S. Jacques, J. Huang, J. Messens, F. Van Breusegem, Oxidative post-translational modifications of cysteine residues in plant signal transduction, J. Exp. Bot. 66 (2015) 2923–2934.

[53] Y. Osakabe, K. Yamaguchi-Shinozaki, K. Shinozaki, L.P. Tran, ABA control of plant macroelement membrane transport systems in response to water deficit and high salinity, New Phytol. 202 (2014) 35–49.

[54] P. Tavladoraki, A. Cona, R. Federico, G. Tempera, N. Viceconte, S. Saccoccio, V. Battaglia, A. Toninello, E. Agostinelli, Polyamine catabolism: target for antiproliferative therapies in animals and stress tolerance strategies in plants, Amino Acids. 42 (2012) 411–426.

[55] T. Van Hautegem, A.J. Waters, J. Goodrich, M.K. Nowack, Only in dying, life: programmed cell death during plant development, Trends Plant Sci. 20 (2015) 102–113.

[56] M. Pireyre, M. Burow, Regulation of MYB and bHLH transcription factors: a glance at the protein level, Mol. Plant. 8 (2015) 378–388.

[57] P.J. Seo, P. Mas, Multiple layers of posttranslational regulation refine circadian clock activity in Arabidopsis, Plant Cell. 26 (2014) 79–87.

[58] R. Stracke, M. Werber, B. Weisshaar, The R2R3-MYB gene family in Arabidopsis thaliana, Curr. Opin. Plant Biol. 4 (2001) 447–456.

[59] G. Mao, X. Meng, Y. Liu, Z. Zheng, Z. Chen, S. Zhang, Phosphorylation of a WRKY transcription factor by two pathogen-responsive MAPKs drives phytoalexin biosynthesis in Arabidopsis, Plant Cell. 23 (2011) 1639–1653.

[60] Z.-J. Wu, X.-H. Li, Z.-W. Liu, H. Li, Y.-X. Wang, J. Zhuang, Transcriptome-wide identification of Camellia sinensis WRKY transcription factors in response to temperature stress, Mol. Genet. Genomics. 291 (2016) 255–269.

[61] A. Taglialatela, S. Alvarez, G. Leuzzi, V. Sannino, L. Ranjha, J.-W. Huang, C. Madubata, R. Anand, B. Levy, R. Rabadan, Restoration of replication fork stability in BRCA1-and BRCA2-deficient cells by inactivation of SNF2-family fork remodelers, Mol. Cell. 68 (2017) 414–430.

[62] A.-L. Jiang, Z.-S. Xu, G.-Y. Zhao, X.-Y. Cui, M. Chen, L.-C. Li, Y.-Z. Ma, Genome-wide analysis of the C3H zinc finger transcription factor family and drought responses of members in Aegilops tauschii, Plant Mol. Biol. Report. 32 (2014) 1241–1256.

[63] C. Yanhui, Y. Xiaoyuan, H. Kun, L. Meihua, L. Jigang, G. Zhaofeng, L. Zhiqiang, Z. Yunfei, W. Xiaoxiao, Q. Xiaoming, The MYB transcription factor superfamily of Arabidopsis: expression analysis and phylogenetic comparison with the rice MYB family, Plant Mol. Biol. 60 (2006) 107–124.

[64] C.S. Silva, S. Puranik, A. Round, M. Brennich, A. Jourdain, F. Parcy, V. Hugouvieux, C. Zubieta, Evolution of the plant reproduction master regulators LFY and the MADS transcription factors: the role of protein structure in the evolutionary development of the flower, Front. Plant Sci. 6 (2016) 1193.

[65] H. Shaked, N. Avivi-Ragolsky, A.A. Levy, Involvement of the Arabidopsis SWI2/SNF2 chromatin remodeling gene family in DNA damage response and recombination, Genetics. 173 (2006) 985–994.

[66] J. Terol, M. Bargues, M. Pérez-Alonso, ZFWD: a novel subfamily of plant proteins containing a C3H zinc finger and seven WD40 repeats, Gene. 260 (2000) 45–53.

[67] R. Hang, C. Liu, A. Ahmad, Y. Zhang, F. Lu, X. Cao, Arabidopsis protein arginine methyltransferase 3 is required for ribosome biogenesis by affecting precursor ribosomal RNA processing, Proc. Natl. Acad. Sci. 111 (2014) 16190–16195.

[68] W.J. Lucas, B.-C. Yoo, F. Kragler, RNA as a long-distance information macromolecule in plants, Nat. Rev. Mol. Cell Biol. 2 (2001) 849.

[69] Q.-H. Wang, C. Zhao, M. Zhang, Y.-Z. Li, Y.-Y. Shen, J.-X. Guo, Transcriptome analysis around the onset of strawberry fruit ripening uncovers an important role of oxidative phosphorylation in ripening, Sci. Rep. 7 (2017) 41477.

[70] W. Xu, C. Dubos, L. Lepiniec, Transcriptional control of flavonoid biosynthesis by MYB–bHLH–WDR complexes, Trends Plant Sci. 20 (2015) 176–185.

[71] X. Li, L.-P. Zhang, L. Zhang, P. Yan, G.J. Ahammed, W.-Y. Han, Methyl Salicylate Enhances Flavonoid Biosynthesis in Tea Leaves by Stimulating the Phenylpropanoid Pathway, Molecules. 24 (2019) 362.

[72] A. Rani, K. Singh, P. Sood, S. Kumar, P.S. Ahuja, p-Coumarate: CoA ligase as a key gene in the yield of catechins in tea [Camellia sinensis (L.) O. Kuntze], Funct. Integr. Genomics. 9 (2009) 271–275.

[73] C.-H. Wang, J. Yu, Y.-X. Cai, P.-P. Zhu, C.-Y. Liu, A.-C. Zhao, R.-H. Lü, M.-J. Li, F.-X. Xu, M.-D. Yu, Characterization and functional analysis of 4-coumarate: CoA ligase genes in mulberry, PLoS One. 11 (2016) e0155814.

[74] X. Zhang, C.-J. Liu, Multifaceted regulations of gateway enzyme phenylalanine ammonia-lyase in the biosynthesis of phenylpropanoids, Mol. Plant. 8 (2015) 17–27.

[75] M.B. Shine, J. Yang, M. El-Habbak, P. Nagyabhyru, D. Fu, D. Navarre, S. Ghabrial, P. Kachroo, A. Kachroo, Cooperative functioning between phenylalanine ammonia lyase and isochorismate synthase activities contributes to salicylic acid biosynthesis in soybean, New Phytol. 212 (2016) 627–636.

[76] Y. Tu, F. Liu, D. Guo, L. Fan, Z. Zhu, Y. Xue, Y. Gao, M. Guo, Molecular characterization of flavanone 3-hydroxylase gene and flavonoid accumulation in two chemotyped safflower lines in response to methyl jasmonate stimulation, BMC Plant Biol. 16 (2016) 132.

[77] S. Miosic, J. Thill, M. Milosevic, C. Gosch, S. Pober, C. Molitor, S. Ejaz, A. Rompel, K. Stich, H. Halbwirth, Dihydroflavonol 4-reductase genes encode enzymes with contrasting substrate specificity and show divergent gene expression profiles in Fragaria species, PLoS One. 9 (2014) e112707.

[78] M.-Z. Nouri, A. Moumeni, S. Komatsu, Abiotic stresses: insight into gene regulation and protein expression in photosynthetic pathways of plants, Int. J. Mol. Sci. 16 (2015) 20392–20416.

[79] P.R. Steed, K.A. Kraft, R.H. Fillingame, Interacting cytoplasmic loops of subunits a and c of Escherichia coli F1F0 ATP synthase gate H+ transport to the cytoplasm, Proc. Natl. Acad. Sci. 111 (2014) 16730–16735.

[80] I. Yruela, Transition metals in plant photosynthesis, Metallomics. 5 (2013) 1090–1109.

[81] M. Weigel, C. Varotto, P. Pesaresi, G. Finazzi, F. Rappaport, F. Salamini, D. Leister, Plastocyanin is indispensable for photosynthetic electron flow in Arabidopsis thaliana, J. Biol. Chem. 278 (2003) 31286–31289.

[82] A. Moolna, C.G. Bowsher, The physiological importance of photosynthetic ferredoxin NADP+ oxidoreductase (FNR) isoforms in wheat, J. Exp. Bot. 61 (2010) 2669–2681.

